# A lipophilicity-based energy function for membrane-protein modelling and design

**DOI:** 10.1101/615658

**Authors:** Jonathan Yaacov Weinstein, Assaf Elazar, Sarel Jacob Fleishman

**Affiliations:** Department of Biomolecular Sciences, Weizmann Institute of Science, Rehovot, Israel

**Keywords:** Rosetta, membrane-protein energetics, *ab initio* structure prediction, *de novo* design, dsTβL, mutational analysis

## Abstract

Membrane-protein design is an exciting and increasingly successful research area which has led to landmarks including the design of stable and accurate membrane-integral proteins based on coiled-coil motifs. Design of topologically more complex proteins, such as most receptors, channels, and transporters, however, demands an energy function that balances contributions from intra-protein contacts and protein-membrane interactions. Recent advances in water-soluble all-atom energy functions have increased the accuracy in structure-prediction benchmarks. The plasma membrane, however, imposes different physical constraints on protein solvation. To understand these constraints, we recently developed a high-throughput experimental screen, called dsTβL, and inferred apparent insertion energies for each amino acid at dozens of positions across the bacterial plasma membrane. Here, we express these profiles as lipophilicity energy terms in Rosetta and demonstrate that the new energy function outperforms previous ones in modelling and design benchmarks. Rosetta ab initio simulations starting from an extended chain recapitulate two-thirds of the experimentally determined structures of membrane-spanning homo-oligomers with <2.5 Å root-mean-square deviation within the top-predicted five models. Furthermore, in two sequence-design benchmarks, the energy function improves discrimination of stabilizing point mutations and recapitulates natural membrane-protein sequences of known structure, thereby recommending this new energy function for membrane-protein modelling and design.

## Introduction

Membrane proteins have essential biological roles as receptors, channels, and transporters. Over the past decade, significant progress has been made in membrane-protein design, including the first design of membrane-integral inhibitors^1^, a transporter^2^, and a *de novo* designed structure based on coiled-coil motifs^3^. Despite this exciting progress, modelling, design, and engineering of membrane proteins lag far behind those of soluble proteins. This lag is due, in part, to the relatively small number of high-resolution membrane-protein structures^4^ and is exacerbated by these proteins’ typically large size. Clearly, however, the most significant complication is that membrane proteins are solvated in a physically heterogeneous and only partly understood environment, comprising water, lipid, and polar lipid headgroups^5^. Modelling solvation is, therefore, a fundamental problem that impacts all membrane-protein structure prediction and design.

Current energy functions used in modelling and design incorporate simplified solvation models^6^. For instance, RosettaMembrane uses information inferred from water-to-hexane partitioning^7^ as a proxy for amino acid solvation in the plasma membrane^8–10^. Due to these simplifications, expert analysis has been a prerequisite for accurate membrane-protein modelling and design^11,12^. Automating modelling and design processes and extending them to complex membrane proteins will likely require an accurate energy function that correctly balances intra-protein interactions, membrane solvation and water solvation^13,14^.

To understand the contributions to membrane-protein solvation, we recently established a high-throughput experimental screen, called deep sequencing TOXCAT-β-lactamase (dsTβL), which quantified apparent amino acid transfer energies from the cytosol to the *E. coli* plasma membrane^15^. From the resulting data, we inferred apparent position-specific insertion profiles for each amino acid relative to alanine, reconciling previously conflicting lines of evidence^16^. Foremost, the lipophilicity inferred for hydrophobic residues, such as Leu, Ile, and Phe, was greater than previously measured in some membrane mimics, including the water-to-hexane transfer energies that are the basis for membrane solvation in Rosetta^7–9^ (approximately 2 kcal/mol according to dsTβL compared to ½ kcal/mol), and in line with theoretical considerations^17,18^. Second, the profiles exhibited a strong 2 kcal/mol preference for Arg and Lys in the intracellular side of the plasma membrane compared to the extracellular side. While this preference, known as the “positive-inside” rule, was revealed based on sequence analysis 30 years ago^19–21^, the dsTβL assay was the first to indicate a large energy gap favouring positively charged residues in the intracellular relative to the extracellular membrane leaflet. The accuracy and generality of the dsTβL apparent transfer energies were partly verified by demonstrating that they correctly predicted the locations and orientations of membrane spans directly from sequence even in several large and complex eukaryotic transporters^22^. Taken together, these results provided reassurance that the dsTβL apparent insertion energies correctly balanced essential aspects of membrane-protein solvation.

As the next step towards accurate all-atom membrane-protein modelling and design, we develop a new lipophilicity-based energy term based on the dsTβL amino acid specific insertion profiles and integrate this energy term in the Rosetta centroid-level and all-atom potentials. We furthermore develop a strategy to enhance conformational sampling of membrane-spanning helical segments and of helix-tilt angles observed in naturally occurring membrane proteins. Encouragingly, the new energy function outperforms previous ones in three benchmarks essential to modelling and design: atomistic *ab initio* structure prediction starting from completely extended chains of single-spanning membrane homo-oligomers of known structure, prediction of mutational effects on stability, and sequence recovery in combinatorial sequence design. Therefore, the combination of lipophilicity and energetics developed for soluble proteins provides a basis for accurate structure prediction and design of membrane proteins.

## Results

### A lipophilicity-based membrane-protein energy function

The recent all-atom energy function in Rosetta, ref2015, is dominated by physics-based terms, including van der Waals packing, hydrogen bonding, electrostatics and water solvation^23^. This energy function was parameterized on a large set of crystallographic structures and experimental data of water-soluble proteins and was shown to outperform previous energy functions in several structure-prediction benchmarks. For membrane-protein modelling and design, however, the ref2015 solvation potential is relevant only to the water-embedded regions of the protein; a different potential is required to model the energetics of amino acids near and within different regions of the plasma membrane.

Accordingly, we sought to replace the ref2015 solvation model with one that encodes a gradual transition from the default water-solvation that evaluates regions distant from the plasma membrane and the dsTβL insertion profiles near and within the plasma membrane. The dsTβL profiles were inferred from an experimental mutation analysis of a monomeric membrane span into which each of the 20 amino acids were individually introduced at each position^15^; the profiles were then normalized to express the apparent transfer energy for each amino acid at each position relative to a theoretical poly-Ala membrane span, yielding apparent ΔΔ*G*_Ala—>mut_ at each position across the plasma membrane (Fig. 1). As a first step to encoding these energy profiles in Rosetta, we smoothed these profiles and symmetrised them with respect to the presumed membrane midplane, except the profiles for Arg, His, and Lys, for which the “positive-inside” rule applies (Supplemental Figure S1).

**Figure 1.**
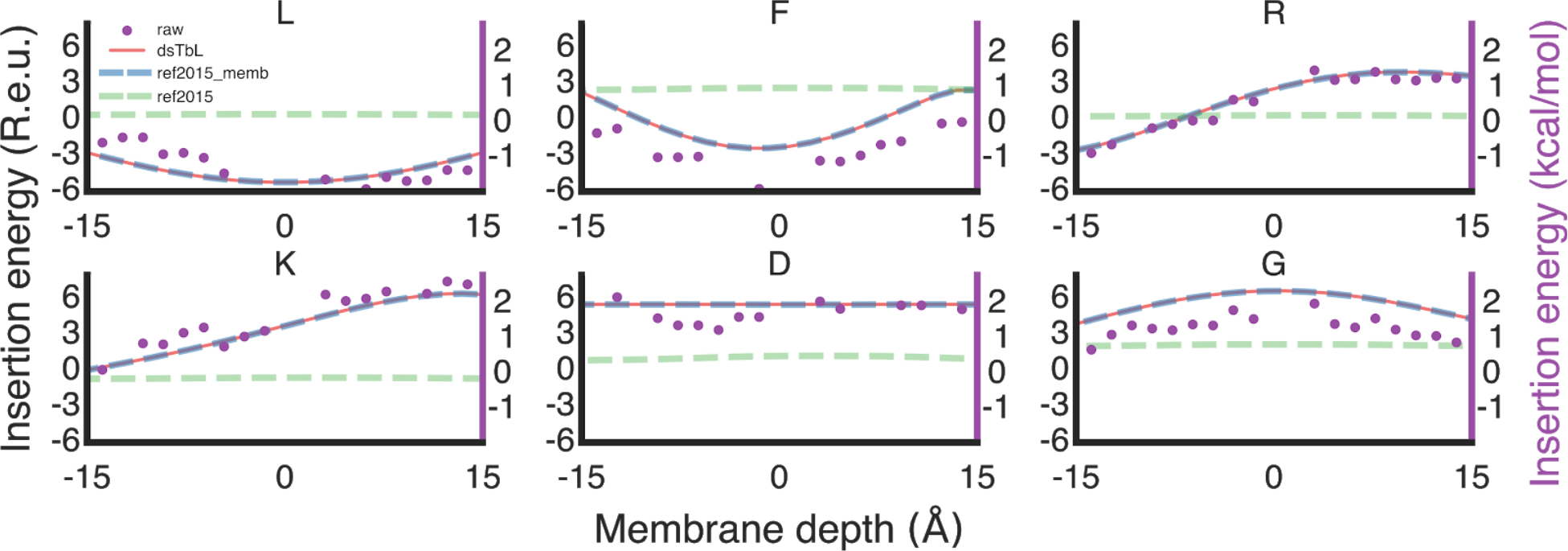
The lipophilicity-based ref2015_memb energy function. Membrane-insertion profiles for six representative amino acids are shown. Raw dsTβL data (purple dots), ref2015 (dashed green line), the ref2015_memb potential (dashed blue line) and the dsTβL profiles (red line). Negative and positive membrane depths indicate the inner and outer membrane leaflets, respectively; the presumed membrane midplane is at 0.

Next, we implemented an iterative strategy to encode the dsTβL energetics in a modified ref2015 all-atom energy function which we called ref2015_memb. To enable efficient conformational search as required in *ab initio* structure prediction and *de novo* design, we also encoded this energetics in the centroid-level energy function^24^. As a reference state in both all-atom and centroid-level modelling, we generated an ideal poly-Ala α helix and placed it perpendicular to the membrane plane. At each position along the helix (including the aqueous and membrane phases), we introduced each of the 19 point mutations, relaxed the models using the all-atom or centroid-level energy functions, and computed the energy difference due to each single-point mutation ΔΔ*G*_Ala—>mut_. In the first iteration of these calculations, the unmodified ref2015 or centroid-level energy functions were used, resulting, as expected, in large deviations from the apparent energies observed in the dsTβL profiles (dashed green lines in Figure 1). We then added a new term, called MPResidueLipophilicity, which encoded the difference between the computed and dsTβL energies for each mutation at each position, ΔΔΔ*G*_Ala—>mut_. We iterated mutation, relaxation, energy calculations, and MPResidueLipophilicity updates for each of the mutations at each position up to ten times, noting that the computed energies converged with the trends observed in the experiment (blue and red lines in Figure 1, respectively). Scripts for calibrating the all-atom and centroid energy functions are available in the supplement to enable adapting future improvements of the Rosetta energy functions to encode the dsTβL energetics.

The dsTβL apparent energy profiles were inferred from a monomeric segment^15^. Consequently, the profiles express the lipophilicity of each amino acid relative to Ala across the membrane when that amino acid is maximally solvent-exposed. To account for amino acid burial in multispan or oligomeric membrane proteins, we derived a continuous, differentiable and easily computable weighting term that expresses the extent of a residue’s burial in other protein segments. For any given amino acid, this weighting term is based on the number of heavy-atom neighbours within 6 and 12 Å distance of the amino acid’s Cβ atom (Eqs. 2–4) resulting in a weight that expresses the extent to which a residue is buried in other protein segments or exposed to solvent (0 to 1, respectively). Water-embedded and completely buried positions are treated with the ref2015 solvation energy; fully membrane-exposed positions are treated with the MPResidueLipophilicity energy, and positions of intermediate exposure are treated with a linearly weighted sum of the two terms.

In summary, the actual contribution from solvation of an amino acid is a function of its exposure to the membrane and depends on the amino acid’s lipophilicity according to the dsTβL apparent energy and the position’s location relative to the membrane midplane. Note that this energy term averages lipophilicity contributions in the plasma membrane and does not express atomic contributions to solvation that are likely to be important in calculating membrane-protein energetics in different types of biological membranes^9,25^, in non-helical membrane-exposed segments, or surrounding water-filled cavities^26^.

The dsTβL assay reports on residue-specific insertion into the plasma membrane. *Ab initio* modelling and *de novo* design, however, also require a potential that addresses the protein backbone solvation. Although the low-dielectric environment in the core of the membrane enforces a strong tendency for forming canonical α helices^5^, deviations from canonical α helicity can make important contributions to membrane-protein structure and function^27^. We, therefore, encoded an energy term, called MPHelicality, that allows sampling backbone dihedral angles and penalises deviations from α helicity (Eq. 5). MPHelicality enforces strong constraints on the dihedral angles in the lipid-exposed surfaces at the core of the membrane and is attenuated in regions that are buried in other protein segments and in the extra-membrane environment (using the same weighting as for lipophilicity, Eq. 1); this term thus allows significant deviations from α helicity only in buried or water-embedded regions.

In preliminary *ab initio* calculations starting from a fully extended chain, we noticed that conformational sampling significantly favoured large helical tilt angles relative to the membrane normal (Θ in Figure 2). By contrast, 50% of naturally observed membrane spans exhibit small tilt angles in the range 15-30°. The skew in conformational sampling towards large tilt angles is expected from previous theoretical investigations according to which the distribution of helix-tilt angles in random sampling is proportional to sin(Θ), substantially preferring large angles compared to the distribution observed in natural membrane proteins^28^. To eliminate this skew in conformational sampling, we introduced another energy term, called MPSpanAngle (Eq. 4 and Fig. 2), that strongly penalized large tilt angles, guiding *ab initio* sampling to tilt angles observed in natural proteins.

**Figure 2.**
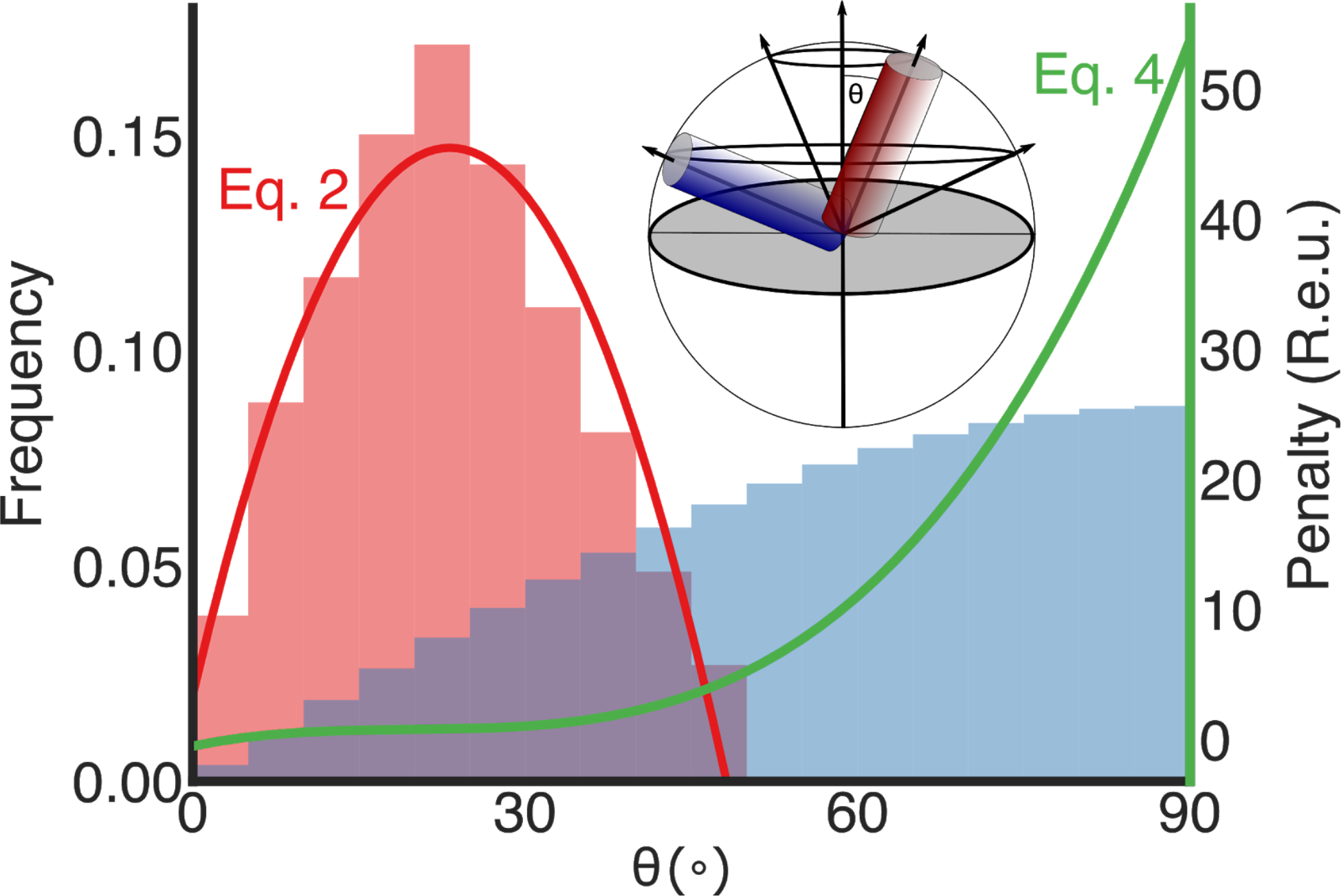
Observed *versus* expected tilt angles in membrane-spanning helices relative to the membrane normal. The distribution of helix tilt angles (Θ in the inset sphere) in natural membrane proteins shows a strong preference for small angles (red bars, left), whereas the distribution resulting from random conformational sampling is proportional to sin(Θ) (blue bars)^28^ significantly overrepresenting large tilt angles. The MPSpanAngle energy term (green line; Eq. 4) penalises large tilt angles and focuses *ab initio* conformational sampling on tilt angles observed in membrane-protein structures. *inset* The expected distribution of helix-tilt angles is proportional to the circumference of a circle plotted by that helix around an axis perpendicular to the membrane-normal (panel adapted from ref. ^28^). The membrane plane is depicted as a grey circle.

In summary, ref2015_memb encodes three new energy terms relative to the soluble energy function ref2015: (1) a lipophilicity term based on amino acid type, membrane-depth, and burial; (2) a penalty on deviations from α helicity in backbone-dihedral angles; and (3) a penalty on the sampling of large tilt angles with respect to the membrane-normal (Supplemental Table 1). In the calculations reported below, the penalties on deviations from α helicity and helix-tilt angles are implemented in all centroid-level *ab initio* structure prediction simulations; all-atom calculations use the ref2015 energy modified with the lipophilicity term.

### *Ab initio* structure prediction in membrane proteins

Previous structure-prediction benchmarks started from canonical α helices or from monomers obtained from experimental structures of homodimers and used the bound-structures in grid search or rigid-body docking^8,9,29–32^. Additionally, structure-prediction studies used experimental constraints, conservation analysis or correlated-mutation analysis to predict residue contacts in order to constrain conformational sampling^11,12,33–38^. Several automated predictors dedicated to single-span homodimers used shape complementarity^39,40^, sequence-packing motifs^41^ or comparative modelling^42^, but to the best of our knowledge, *ab initio* modelling calculations, starting from a fully extended chain, have not been described. Given that deviations from canonical α helicity make important contributions to membrane-protein structure and function^27^, we decided to apply a more stringent test using *ab initio* modelling, sampling all symmetric backbone, sidechain, and rigid-body degrees of freedom.

To test *ab initio* modelling using the new energy function, we applied the fold-and-dock protocol^43^, which has been successfully applied in a variety of soluble-protein structure prediction and design studies^44–47^. Briefly, fold-and-dock starts from an extended chain and conducts several hundred iterations of symmetric centroid-level backbone-fragment insertion and relaxation moves. It then applies symmetric all-atom refinement including all dihedral sidechain and backbone degrees of freedom (Supplemental Movie 1). To generate an energy landscape, we ran 5,000 independent trajectories (50,000 for high-order oligomers) for every 19 and 21 residue subsequence of each homooligomer, filtered the resulting models according to energy and structure parameters (Methods), and isolated the lowest-energy 10% of the models. Models were then clustered according to their energies and conformations, and five cluster representatives were compared to the experimental structures (Figures 2 and 3, Table 1). For comparison, we applied the described methodology using ref2015_memb, ref2015 and the current membrane-protein energy function in Rosetta,RosettaMembrane^8–10^.

**Figure 3.**
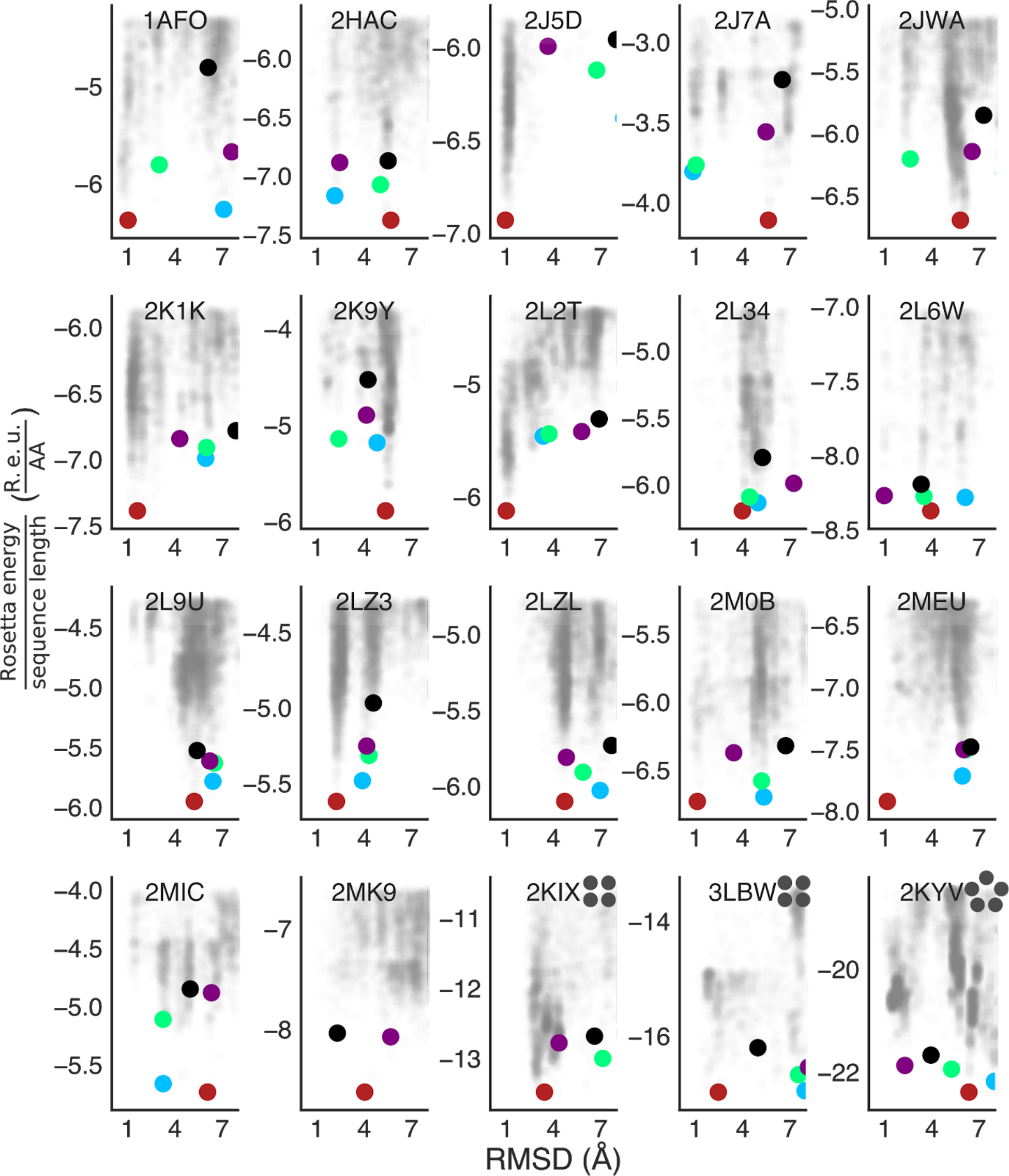
Energy landscapes for the *ab initio* structure prediction benchmark. All models that passed the energy and structure-based filters are shown as semi-transparent grey dots. Each of the five lowest-energy clusters is indicated by coloured circles. The PDB entry is indicated on each panel and the oligomeric state is specified by grey circles for higher oligomeric states than homodimers. Y-axes report the ref2015_memb energy normalised by the monomeric sequence length of each model.

**Table 1.** Structure prediction benchmark. Grey cells indicate RMSD < 2.5Å or fraction of native contacts using ref2015_memb > 0.7.

| PDB<br>code | # subunits<sup>1</sup> | fraction of<br>native<br>contacts | RMSD of nearest model structure (Å) |  |  |  |  |
| --- | --- | --- | --- | --- | --- | --- | --- |
|  |  |  | ref2015_memb | RosettaMembrane | ref2015 | PREDDIMER | TMDIM |
| 2J5D | 2 | 0.92 | 0.95 | 1.04 | 1.06 | 8.37 | 2.44 |
| 2L6W | 2 | 0.90 | 0.98 | 2.67 | 2.04 | 2.28 | 5.22 |
| 1AFO | 2 | 0.87 | 1.02 | 2.87 | 1.13 | 1.99 | 0.84 |
| 2L2T | 2 | 0.86 | 1.01 | 4.77 | 6.40 | 1.85 | 0.75 |
| 2MEU | 2 | 0.80 | 1.17 | 2.81 | 4.65 | 2.71 | 4.05 |
| 2J7A | 2 | 0.80 | 0.85 | 3.36 | 0.94 | 9.74 | 6.54 |
| 2M0B | 2 | 0.72 | 1.12 | 1.57 | 1.55 | 3.22 | 1.78 |
| 2K9Y | 2 | 0.58 | 2.39 | 1.82 | 1.59 | 1.89 | 3.88 |
| 2K1K | 2 | 0.54 | 1.62 | 1.38 | 1.18 | 1.77 | 1.51 |
| 2LZ3 | 2 | 0.47 | 2.24 | 2.14 | 1.95 | 2.09 | 3.56 |
| 2MK9 | 2 | 0.41 | 2.30 | 2.15 | 2.56 | 6.88 | 2.95 |
| 2HAC | 2 | 0.30 | 2.13 | 3.24 | 2.31 | 2.06 | 2.34 |
| 2JWA | 2 | 0.17 | 2.63 | 1.36 | NA | 2.42 | 2.21 |
| 2LZL | 2 | 0.00 | 4.72 | 4.54 | NA | 3.64 | 3.55 |
| 2L9U | 2 | 0.00 | 5.22 | 4.15 | NA | 4.24 | 1.66 |
| 2L34 | 2 | 0.00 | 3.98 | 3.92 | NA | 1.12 | 4.84 |
| 2MIC | 2 | 0.00 | 3.25 | 3.33 | 4.86 | 8.70 | 5.57 |
| 3LBW | 4 | 0.14 | 2.45 | 1.82 | 7.10 | NA | NA |
| 2KIX | 4 | 0.10 | 3.43 | 4.06 | NA | NA | NA |
| 2KYV | 5 | 0.22 | 2.29 | 1.82 | 1.46 | NA | NA |

<sup>1</sup> oligomeric state (dimer, tetramer, or pentamer)

The Protein Data Bank (PDB) contains 17 nonredundant (sequence identity <80%) NMR and X-ray crystallographic structures of natural single-span homodimers, two tetramers and one pentameric structure. Of the 20 cases in the benchmark, fold-and-dock simulations using ref2015_memb predicted near-native (<2.5 Å root-mean-square deviation [RMSD]) low-energy models for 14 homooligomers compared to nine using RosettaMembrane; the soluble energy function ref2015 also resulted in nine correct predictions. Moreover, prediction rates using ref2015_memb were similar for left-as for right-handed homodimers (Supplemental Table S2) and in 11 cases, the top 3 lowest-energy predicted models contained a near-native prediction (Fig. 3). Of the three high-order oligomers tested, ref2015_memb successfully recapitulated the structures of the M2 tetramer and phospholamban pentamer. The PREDDIMER^40^ and TMDIM^41^ structure-prediction web servers, which do not use *ab initio* modelling, found models at <2.5 Å RMSD for nine and eight of the 17 homodimers, respectively. Thus, *ab initio* calculations using ref2015_memb accurately predict structures in two-thirds of the homooligomers in our benchmark, including high-order oligomers that cannot be predicted by other automated methods.

The successfully predicted homooligomers exhibit different structural packing motifs. The majority of the homodimer interfaces are mediated by the ubiquitous Gly-xxx-Gly motif^48^, in which two small amino acids separated by four positions on the primary sequence enable close packing between the helices. There is uncertainty whether these motifs additionally form stabilising Cα hydrogen bonds^49,50^. Our structure-prediction analysis cannot resolve this uncertainty; note, however, that the new energy function ref2015_memb does not encode terms for Cα hydrogen bonds and yet recapitulates a large fraction of the homodimer structures (Figures 3 and 4, and Table 1). The underlying reason for successful prediction is that the dsTβL energetics encodes a strong penalty on exposing Gly residues to the lipid bilayer (approximately 2 kcal/mol/Gly at the membrane mid-plane; Figure 1), driving the burial of Gly amino acids within the homodimer interface (*i.e.*, “solvophobicity”). Thus, lipophilicity and interfacial residue packing are sufficient for accurate structure prediction in a large fraction of the targets we examined.

**Figure 4.**
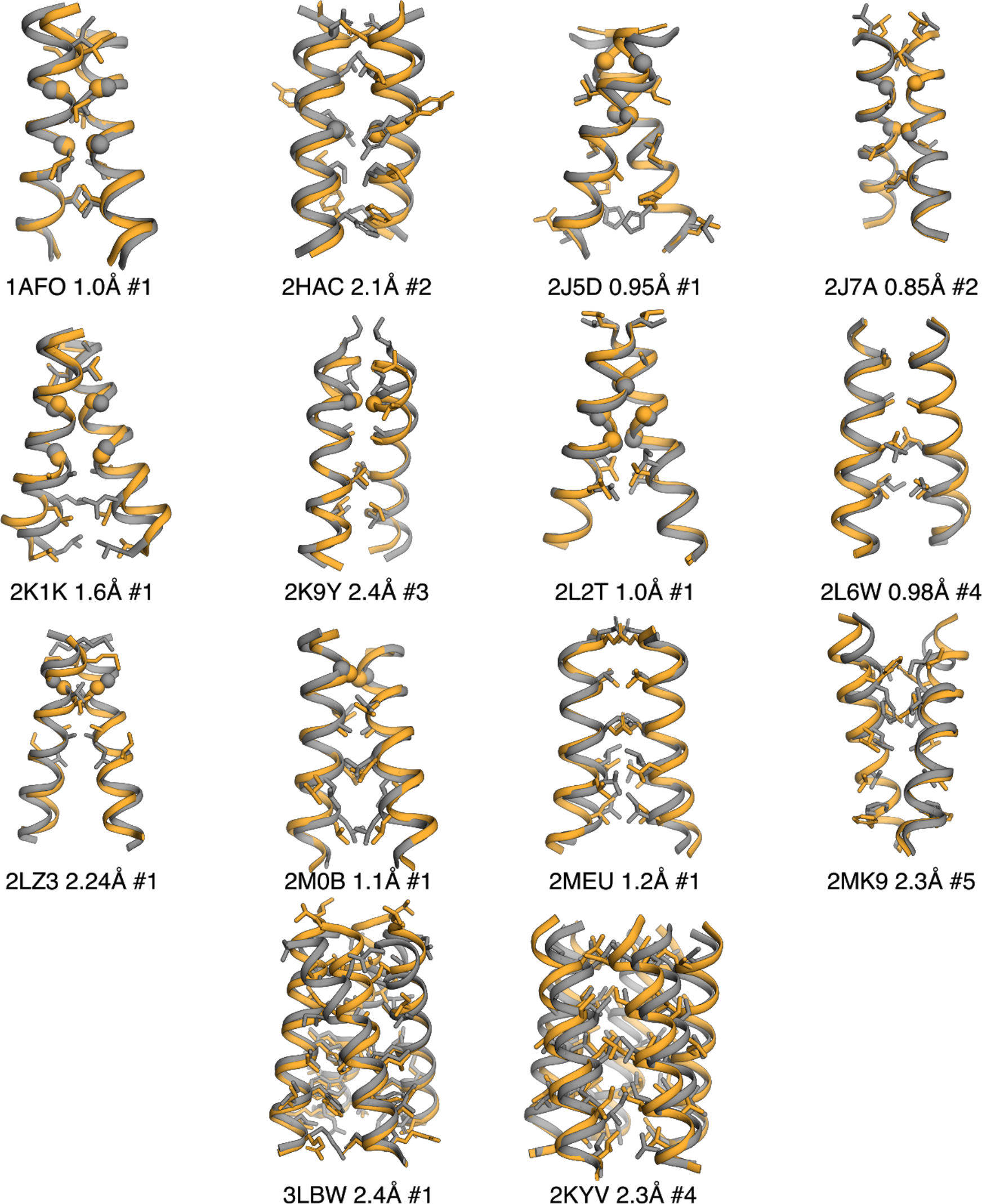
Structural comparison of the top-predicted model (by RMSD) to the experimentally determined structure. PDB entry, RMSD and the model’s ranking (in energy) among the top-5 predicted models are indicated. Only accurately predicted structures (< 2.5 Å) are shown.

Using the dsTβL assay, we also examined the effects of dozens of point mutations in glycophorin A on apparent association energy (ΔΔ*G*_*binding*_) in the bacterial plasma membrane^15^. As a stringent test of the new energy function, we conducted fold-and-dock calculations using both ref2015_memb and RosettaMembrane starting from the sequences of each of the point mutants. To reduce uncertainty in interpreting the experimental results, we focused on 32 mutations that exhibited large apparent energy changes in the experiment (|ΔΔ*G*_*binding*_| ≥ 2 kcal/mol) and compared the median computed ΔΔ*G*_*binding*_ of the lowest-energy models to the experimental observation (Fig. 5, Supplemental Table S3). ref2015_memb outperformed RosettaMembrane, correctly assigning 81% of mutations as stabilizing or destabilizing compared to 66% for RosettaMembrane. Note that as observed in studies of mutational effects on stability in soluble proteins, the correlation coefficient between computed and observed values was low (Pearson *r*^2^=0.21 and 0.02 for ref2015_memb and RosettaMembrane, respectively)^51–54^. Such low correlation coefficients provide an impetus for improving the energy function; however, as we previously demonstrated, discriminating stabilizing from destabilizing mutations is sufficient to enable the design of accurate, stable, and functionally efficient proteins^54–59^.

**Figure 5.**
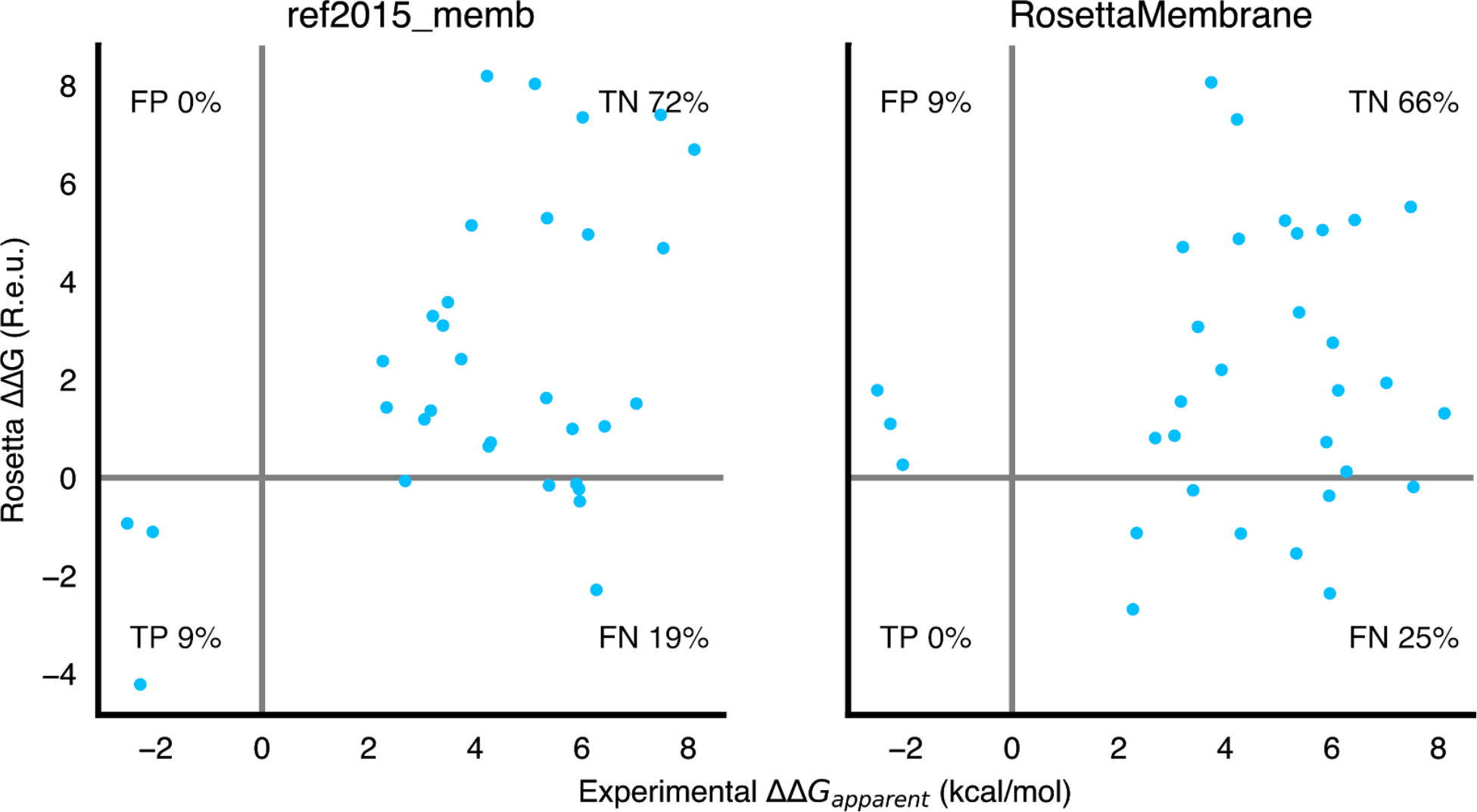
Predicted versus experimental ΔΔ*G*_*binding*_ values of single-point mutations in glycophorin. **A.** The structure of every point mutant was predicted *ab initio*, and the median ΔΔ*G*_*binding*_ relative to the wild type sequence is reported. Only point mutations that exhibited |ΔΔ*G*_*binding*_| ≥ 2 kcal/mol in the experiment were analysed. TP, TN, FP, and FN — true positive, true negative, false positive, and false negative, respectively.

We next tested sequence-recovery rates using combinatorial sequence optimisation based on ref2015, ref2015_memb, and RosettaMembrane in a benchmark of 20 non-redundant structures (<80% sequence identity) ranging in size from 124-765 amino acids^60^. ref2015_memb outperformed the other energy functions, exhibiting 83% sequence recovery, on average, when each design was compared to the target’s natural homologs (Table 2). To our surprise, the soluble energy function ref2015 outperformed RosettaMembrane in this test and was almost as successful as ref2015_memb (78% overall success), implying that the packing and electrostatic models of ref2015^23^ enabled at least some of the improvement observed in sequence recovery by ref2015_memb (see Supplemental Table 1 for a comparison of the energy functions). High sequence recovery in both buried and exposed positions implies that ref2015_memb may be applied effectively to design large and complex membrane proteins.

**Table 2.** Sequence recovery rates in Rosetta combinatorial sequence optimisation.

|  | Sequence recovery<sup>1</sup> |  |  | Homology recovery<sup>2</sup> |  |  |
| --- | --- | --- | --- | --- | --- | --- |
|  | buried | exposed | all | buried | exposed | all |
| ref2015_memb | 0.52 | 0.32 | 0.42 | 0.86 | 0.81 | 0.83 |
| ref2015 | 0.53 | 0.33 | 0.43 | 0.85 | 0.71 | 0.78 |
| RosettaMembrane | 0.23 | 0.20 | 0.21 | 0.64 | 0.70 | 0.67 |

<sup>1</sup> Only exact matches to the natural protein sequence are counted as recovered
<sup>2</sup> For each target protein, a position-specific scoring matrix (PSSM) was computed from a multiple-sequence alignment. At each position, recovery was considered if the amino acid identity had a PSSM score $\geq 0$ .

## Discussion

An accurate energy function is a prerequisite for automated modelling and design, and solvation makes a critical contribution to protein structure and function. The recent dsTβL apparent energies of insertion into the plasma membrane^15^ enabled us to derive an empirical lipophilicity-based energy function for Rosetta. The results demonstrate that ref2015_memb outperforms RosettaMembrane in three benchmarks that are important for structure prediction and design. As ref2015_memb is based on the current state-of-the-art water-soluble Rosetta energy function, prediction accuracy is high for ref2015_memb both in soluble regions and in the core of the membrane domain. Thus, the lipophilicity preferences inferred from the dsTβL energetics together with the residue packing calculations in Rosetta enable accurate modelling in several *ab initio* prediction cases. The current energy function and the fold-and-dock procedure accurately model homooligomeric interactions in the membrane and the effects of point mutations, suggesting that they may enable the accurate design of homooligomeric single-span receptor-like transmembrane domains.

Nevertheless, certain important attributes of membrane-protein energetics are not yet addressed by ref2015_memb; for instance, atomic-level solvation and the impact on electrostatic interactions due to changes in the dielectric constant in various parts of the membrane are currently not treated^8,26^ and warrant further research. The benchmark reported here provides a basis on which improvements in the energy function can be verified.

We recently showed that evolution-guided atomistic design calculations, which use phylogenetic analysis to guide atomistic design calculations^61^, enabled the automated, accurate and effective design of large and topologically complex soluble proteins. Designed proteins exhibited atomic accuracy, high expression levels, stability^54,55^, binding affinity, specificity^59^, and catalytic efficiency^57,58^. Membrane proteins are typically large and challenging targets for conventional protein-engineering and design methods. Looking ahead, we anticipate that evolution-guided atomistic design using the improved energy function may enable reliable design in this important but often formidable class of proteins.

## Methods

### Rosetta source code

All code is available in the Rosetta release at www.rosettacommons.org. Command lines and RosettaScripts^62^ are available in the supplement.

### Membrane-insertion profiles

The original dsTβL insertion profiles^15^ were modified to generate smooth and symmetric functions^22^. The polar and charged residues Asp, Glu, Gln and Asn, which exhibited few counts in the deep sequencing analysis, were averaged such that the insertion energy at the membrane core (−10 to 10 Å; negative values correspond to the inner membrane leaflet and positive values to the outer leaflet) was applied uniformly to the entire membrane span. The profile for His was capped at the maximal value observed in the experiment (2.3 kcal/mol) between 0Å (membrane midplane) and 20 Å. The dsTβL profile for Cys is unusually asymmetric. Cys residues are rare in membrane proteins^63^ and are likely to have similar polarity to Ser. We, therefore, applied the profile measured for Ser to Cys. To convert the values from the dsTβL insertion profiles to Rosetta energy units (R.e.u.) they were multiplied by 2.94 following interpolation reported in ref. ^23^. The dsTβL profiles spanned 27 positions, and we correspondingly translated them to span −20 to +20 Å relative to the membrane midplane.

### Residue lipophilicity

The context-dependent, one-body energy term MPResidueLipophilicity was implemented to encode the dsTβL insertion profiles in ref2015. Starting from an ideal poly Ala α helix embedded perpendicular to a virtual membrane, every position was mutated to all 19 identities, relaxed, and the energy difference between the ref2015 energy and the dsTβL energy was implemented in MPResidueLipophilicity. This process was repeated ten times to reach convergence, and the resulting energy profiles were fitted by a cubic spline^64^, generating continuous, differentiable functions for all 19 amino acids relative to Ala, which was assumed to be 0 throughout the membrane. The splines were recorded in the Rosetta database and are loaded at runtime. Insertion profiles adjustments were done using a python3 script available at github.com/Fleishman-Lab/membrane_protein_energy_function.

### Residue burial

The number of protein atoms within 6 and 12 Å of each amino acid’s Cβ atom is computed and transformed to a burial score (Eq. 1). We used sigmoid functions which range from 0 to 1, corresponding to completely lipid-exposed and completely buried,respectively.

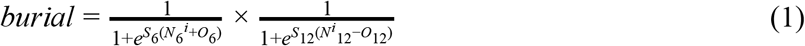

Where *N* is the number of heavy atoms and *S* and *O* determine the slope and offset of the sigmoids and are different for all-atom and centroid calculations. Each parameter has different thresholds at 6 or 12 Å. For all-atom calculations, *S* = 0.15 and 0.5 and *O* = 20 and 475, for 6 and 12 Å radii, respectively. For centroid-level calculations, *S* = 0.15 and 5 and O = 20 and 220 for 6 and 12Å radii, respectively. For each amino acid, the product of the 6 and 12Å sigmoid functions is taken, producing a continuous, differentiable function that transitions from buried to exposed states. These parameters were determined by visualising the burial scores of all amino acids in several polytopic membrane proteins of known structure.

### Tilt-angle (Θ; Fig. 2) penalty

All membrane-spanning helices reported in the PDBTM^65^ dataset (version 20170210) were analyzed for their tilt angles with respect to the membrane normal. A second-degree polynomial was fitted to this distribution using scikit-learn^66^.

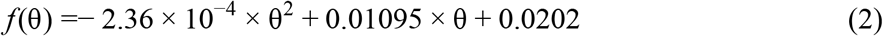

As Bowie noted, the expected distribution function of helix-tilt angles is sin(**Θ**)^28^. We, therefore, used a partition function to convert the expected distribution (sin(**Θ**)) and observed one (Eq. 2) to energy functions, finally subtracting the expected energy from the observed one to derive the helix-tilt penalty function:

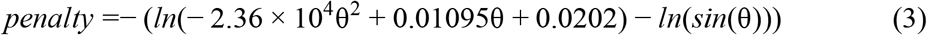

Where θ is given in degrees. In order to simplify runtime calculations, we approximated Eq. 3 using a third-degree polynomial (using scikit-learn) (Fig. 2).

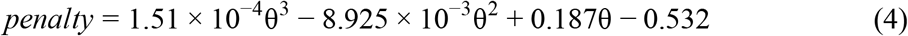

### Penalizing deviations from ideal α helicity

The MPHelicality energy term penalizes the energy of every position that exhibits ϕ-ψ torsion angles significantly different from ideal α helices. A paraboloid function was manually calibrated to express a penalty for any given (ϕ, ψ). The paraboloid centre, for which the penalty is 0, was set to the centre of the helical region according to the Ramachandran plot (ϕ=60°, ψ=45°)^67^. The paraboloid curvature was set to 25, such that the penalty is low throughout the ϕ-ψ torsion angles space observed for α helices^67^. As segments buried against the protein should not be penalized to the same extent as those completely exposed to the membrane, the burial approximation of Equation 1 is used to weight MPHelicality. Moreover, as the protein extends outside of the membrane, the penalty is attenuated with a function that follows the trend observed for the hydrophobic residues, Leu, Ile, and Phe (see Fig. 1A). In effect, the MPHelicality term favours α helicity in lipid-exposed surfaces in the core of the membrane, thereby enforcing some of the electrostatic and solvophobic effects that are essential for correctly modelling the backbone but are not expressed in the residue-specific dsTβL energy profiles.

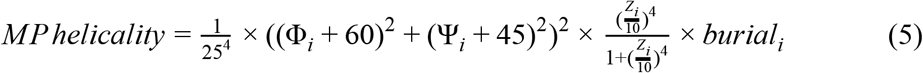

Where ϕ and ψ are given in degrees, *Z* is the distance from the membrane midplane of residue *i*, and burial is calculated as in Eq 1.

### A benchmark for structure prediction of single-span homooligomers

17 structures of single-span homodimers, two homotetramers and one pentamer were selected from the PDB (Supplemental Table 2). For each structure, a 20-30 residue segment comprising the membrane-spanning domain was manually chosen. A sliding window then extracted all 19 or 21 residue subsequences. For each subsequence, three and nine residue backbone fragments were generated using the Rosetta fragment picker application^68^. The fold-and-dock protocol^43^ was used to compute 5000 models (50,000 models for tetramers and the phospholamban pentamer), and the lowest-energy 10% of the models were subsequently filtered using structure and energy-based filters (solvent accessible surface area >500 Å; shape complementarity^69^ *Sc*>0.5; ΔΔG_binding_ <−5 R.e.u.; rotameric binding strain^70^ < 4 R.e.u.; helicality <0.1 R.e.u. (computed using Eq. 5); and closest distance between the interacting helices < 9 Å, as calculated by the filter HelixHelixAngle). For each target, the filtered models from all subsequences were then pooled together and clustered using a score-wise clustering algorithm. This is an iterative process, where each iteration calculates the RMSD of all unclustered models to the best-energy model, and removes the ones closer than 4 Å. RMSD to NMR structures were calculated with respect to the first model in the PDB entry.

### A benchmark for ΔΔ*G*_*binding*_ prediction of single-spanning homodimers

Glycophorin A mutants that exhibited |ΔΔ*G*_*binding*_| > 2 kcal/mol according to the dsTβL study^15^ were modelled using the same fold-and-dock protocol described for the structure prediction of homodimers. The median of computed ΔΔ*G*_*binding*_ for the top models is reported.

### Sequence-recapitulation benchmark

20 structures of polytopic membrane-spanning proteins were taken from ref. ^60^, 11 of which were symmetric complexes^60^. All were refined (eliminating sidechain conformation information before refinement), and for each protein, 100 designs were computed using combinatorial sequence design followed by sidechain and backbone minimization, and the lowest-energy 10 designs were checked for the fraction of mutations relative to the target protein. For each target protein, a multiple-sequence alignment was prepared: homologous sequences were automatically collected using BLASTP^71^ on the nonredundant sequence database^72^ with a maximal number of targets set to 3,000 and an *e*-value ≤ 10^−4^. All sequences were clustered using CD-hit^73^ with a 90% sequence identity threshold. Sequences were then aligned using MUSCLE^74^ with default parameters. A position-specific scoring matrix (PSSM) was calculated using PSI-BLAST^75^. In the sequence-recovery benchmark, where homologous sequences are considered, the substitution of a given position to an identity with a PSSM score ≥ 0 is considered a match.

## Acknowledgements

We thank Rebecca Alford for help with the membrane protein framework in Rosetta and for suggesting the use of splines for fitting the dsTβL profiles. We also thank Adi Goldenzweig, Olga Khersonsky and Saar Shoer for helpful comments. The research was supported by a charitable donation from Sam Switzer and family and from Anne Christopoulos and Carolyn Hewitt.

