## Supplemental information for "A lipophilicity-based energy function for membrane-protein modelling and design"

| Energy term | weight | |
|---|---|---|
| | ref2015_memb | RosettaMembrane |
| fa_atr | 1 | 0.8 |
| fa_dun | 0.7 | 0.01 |
| fa_elec | 1 | 0.026 |
| fa_intra_rep | 0.005 | 0.004 |
| fa_rep | 0.55 | 0.44 |
| fa_sol | 1 | 0 |
| hbond_bb_sc | 1 | 2.34 |
| hbond_lr_bb | 1 | 1.17 |
| hbond_sc | 1 | 2.2 |
| hbond_sr_bb | 1 | 1.17 |
| omega | 0.4 | 0.5 |
| p_aa_pp | 0.6 | 0.32 |
| pro_close | 1.25 | 1 |
| rama_prepro | 0.45 | (rama) 0.2 |
| ref | 1 | 1 |
| dslf_fa13 | 1.25 | |
| dslf_ca_dih | | 5 |
| dslf_cs_ang | | 2 |
| dslf_ss_dih | | 5 |
| dslf_ss_dst | | 0.5 |
| fa_intra_sol_xover4 | 1 | |
| yhh_planarity | 0.625 | |
| lk_ball_wtd | 1 | |
| mp_res_lipo | 1 | |
| mp_span_angle | 1[1] | |
| mp_helicality | 1[2] | |
| fa_mpenv | | 0.3 |
| fa_mpenv_smooth | | 0.5 |
| fa_mpsolv | | 0.35 |
| fa_pair | | 0.49 |

**Supplemental Table 1. Comparison of energy term weights.** ref2015_memb is based on the ref2015 energy function whereas RosettaMembrane is based on score12. For a detailed explanation on energy terms see ref[23,24,76].

[1] centroid level energy functions.

[2] centroid level energy functions, and full-atom energy functions for single spanning proteins.

| PDB | # subunits | Name | handedness | method |
|---|---|---|---|---|
| 2L2T | 2 | ErbB4 | right | NMR |
| 2JWA | 2 | ErbB2 | right | NMR |
| 2MEU | 2 | VEGFR2 mutant | parallel | NMR |
| 2LZL | 2 | FGFR3tm | left | NMR |
| 1AFO | 2 | GpA | right | NMR |
| 2K9Y | 2 | EphA2 | left | NMR |
| 2MK9 | 2 | TLR3 | right | NMR |
| 2HAC | 2 | Zeta-Zeta TM dimer | left | NMR |
| 2LCX | 2 | ErbB4 | right | NMR |
| 2K1K | 2 | EphA1 at pH=4.3 | right | NMR |
| 2J5D | 2 | BNIP3 | right | NMR |
| 2M0B | 2 | ErbB1 | right | NMR |
| 2L9U | 2 | ErbB3 | left | NMR |
| 2L34 | 2 | DAP12 | left | NMR |
| 2L6W | 2 | PDGFR beta-TM | left | NMR |
| 2LZ3 | 2 | amyloid precursor protein | right | NMR |
| 2J7A | 2 | NrfH Cytochrome C Quinol Dehydrogenase | right | X-ray |
| 2MIC | 2 | p75 | right | NMR |
| 2KIX | 4 | Channel domain of BM2 protein from influenza B virus | | NMR |
| 3LBW | 4 | M2 closed state | | X-ray |
| 2KYV | 5 | phospholamban | | NMR |

**Supplemental Table 2. Structures in the prediction benchmark.**

| sequence position | wild-type identity | mutations identity | Experimental ΔΔG (kcal/mol) | RosettaMembrane | ref2015_memb |
|---|---|---|---|---|---|
| 10 | A | F | 3.17 | 1.55 | 1.37 |

| | | | | | |
|---|---|---|---|---|---|
| 10 | A | I | 3.05 | 0.86 | 1.19 |
| 10 | A | L | 3.40 | -0.26 | 3.10 |
| 10 | A | M | 4.29 | -1.14 | 0.72 |
| 11 | G | A | 3.49 | 3.08 | 3.58 |
| 11 | G | F | 7.53 | -0.19 | 4.68 |
| 11 | G | I | 7.48 | 5.52 | 7.40 |
| 11 | G | L | 8.11 | 1.31 | 6.70 |
| 11 | G | M | 7.02 | 1.94 | 1.51 |
| 11 | G | V | 6.43 | 5.26 | 1.05 |
| 14 | G | A | 2.69 | 0.81 | -0.07 |
| 14 | G | F | 5.96 | -2.36 | -0.48 |
| 14 | G | I | 5.90 | 0.73 | -0.13 |
| 14 | G | L | 5.95 | -0.37 | -0.23 |
| 14 | G | M | 6.27 | 0.12 | -2.29 |
| 14 | G | V | 5.39 | 3.37 | -0.16 |
| 7 | G | A | 3.20 | 4.71 | 3.30 |
| 7 | G | F | 3.93 | 2.20 | 5.15 |
| 7 | G | I | 5.12 | 5.25 | 8.03 |
| 7 | G | L | 4.22 | 7.31 | 8.20 |
| 7 | G | M | 5.35 | 4.98 | 5.30 |
| 7 | G | V | 3.74 | 8.07 | 2.42 |
| 17 | L | V | -2.53 | 1.79 | -0.93 |
| 3 | L | A | -2.29 | 1.10 | -4.22 |
| 15 | T | A | 2.34 | -1.13 | 1.43 |
| 15 | T | F | 6.12 | 1.78 | 4.96 |
| 15 | T | I | 5.82 | 5.05 | 1.00 |
| 15 | T | L | 6.02 | 2.75 | 7.35 |
| 15 | T | M | 5.33 | -1.54 | 1.63 |
| 15 | T | V | 4.25 | 4.87 | 0.64 |
| 12 | V | A | -2.05 | 0.27 | -1.11 |
| 12 | V | L | 2.27 | -2.68 | 2.38 |

**Supplemental Table 3. Raw data on the $\Delta\Delta G_{binding}$ prediction data.**

| PDB | Protein Name | # subunits |
|---|---|---|
| 1FX8 | Glycerol Facilitator | Tetramer |
| 1K4C | KcsA Potassium Channel, H+ with Fab | Tetramer |

| | | |
|---|---|---|
| 1M0L | Bacteriorhodopsin | Trimer |
| 1OTS | H+/Cl- exchange transporter | Dimer |
| 1U19 | Rhodopsin (bovine outer segment) | Monomer |
| 2C3E | Mitochondrial ADP/ATP Carrier | Monomer |
| 2UUI | (Apo) Leukotriene Synthase | Trimer |
| 2VPZ | Polysulfide Reductase | Dimer |
| 2XOV | Rhomboid-Family intramembrane protease | Monomer |
| 3B9W | Rh50 protein | Trimer |
| 3GIA | (Apo) ApcT Na+-independent Amino Acid Transporter | Monomer |
| 3K3F | Urea Transporter | Trimer |
| 3KLY | FocA formate transporter w/o formate | Pentamer |
| 3M71 | SLAC1 anion channel TehA homolog | Trimer |
| 3O0R | Nitric Oxide Reductase subunit B | Monomer |
| 3RLB | ThiT, S component of the Thiamin Transporter | Dimer |
| 3V5U | Sodium Calcium Exchanger (MCX) | Monomer |
| 3ZOJ | AQY1 Yeast Aquaporin | Tetramer |
| 4A2N | Isoprenylcysteine carboxyl methyltransferase | Monomer |
| 4IKV | Proton-dependent oligopeptide transporter | Monomer |

**Supplemental Table 4. Structures for the sequence recovery benchmark.**

**Supplemental Movie 1**. A representative fold and dock simulation for glycophorin A. In the first 25 seconds of the movie, simulations are in centroid mode, followed by all-atom refinement.

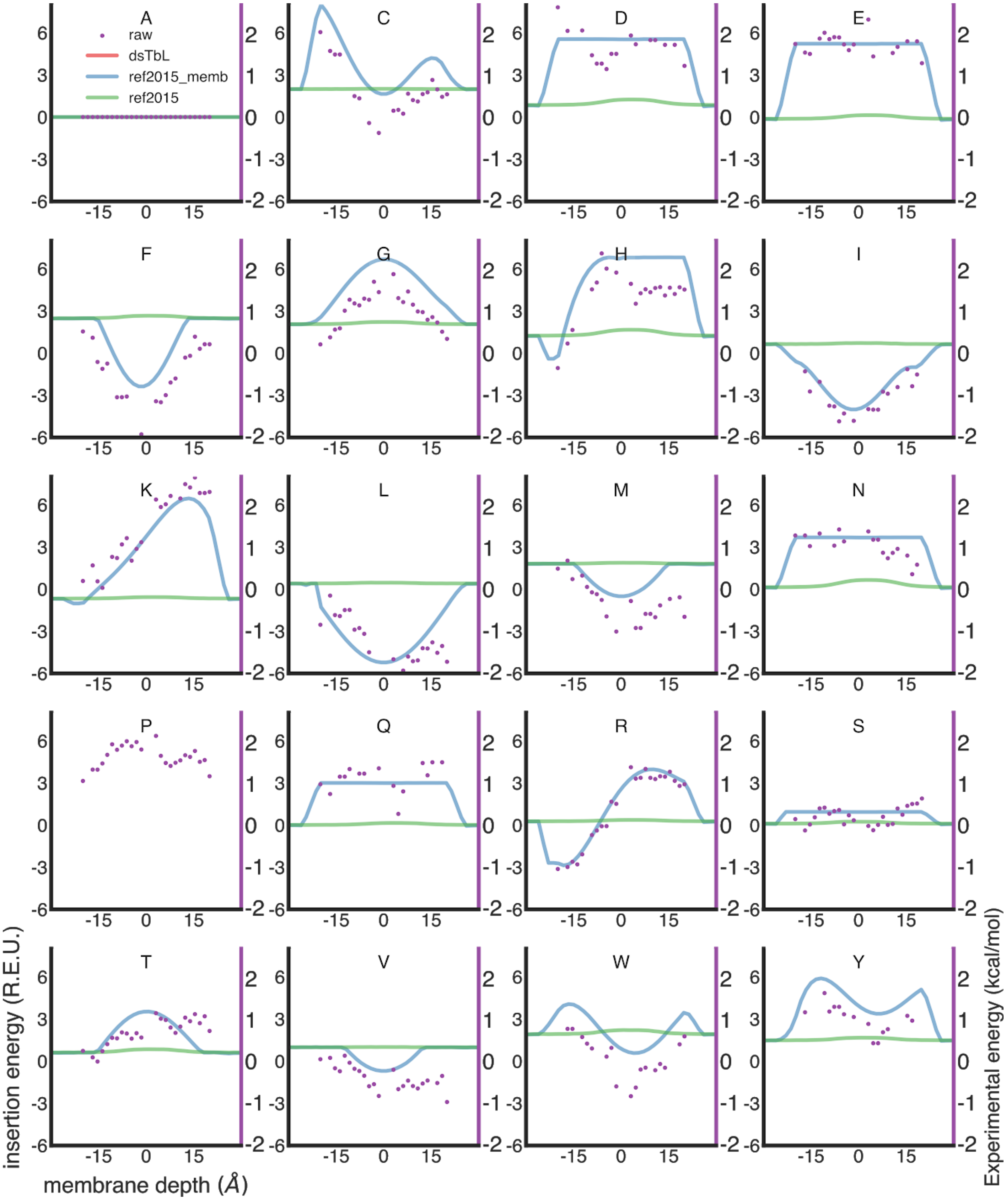


**Figure S1. Adjustment and calibration of insertion profiles. Each panel shows membrane insertion profiles for a different amino acid. Raw dsTβL data (purple dots, right-hand Y-axis),** dsTβL adjusted profiles (red line), ref2015 insertion profiles (green) and ref2015_memb profiles (blues dashed line). Different residues affect the α helix differently, and therefore have different baselines. Note that Pro has no profile under ref2015_memb due to its effect on the backbone.

Fragment files were created for each sequence using the following command line:

```
fragment_picker  -database /Rosetta/main/database
-in::file::vall /Rosetta/tools/fragment_tools/vall.jul19.2011.gz
-in::file::fasta fasta_file -frags::ss_pred ss2_file predA -s
arbiotrary_pdb -frags::scoring::config
fragment_picker_simple.wghts -frags::bounded_protocol
-frags::frag_sizes 3 -frags::n_candidates 200 -frags::n_frags
200 -frags::describe_fragments frags.fsc -frags:allowed_pdb
pdb_chains_in_vall_fragment_picker_12Jul.txt
-out::file::frag_prefix fragment_file
```

Where fragment_picker_simple.wghts is:

```
# score name   priority  wght max_allowed  extras
RamaScore               400     2.0      -        predA
SecondarySimilarity     350     1.0      -        predA
FragmentCrmsd           0       0.0      -
```

In order to use membrane proteins only, a list of membrane protein PDBs is provided to the flags frags:allowed_pdb:

```
1a0tP  1ek9A  1eysC  1eysH  1fepA  1h2sA  1h2sB  1i78A  1j4nA  1jb0A
1jb0B  1jb0C  1jb0D  1jb0E  1jb0F  1jb0J  1jb0K  1jb0L  1kf6A  1kf6B
1kf6C  1kf6D  1kmoA  1kqfA  1kqfB  1kqfC  1ldfA  1lghA  1lghB  1m0kA
1m56C  1m56D  1nkzA  1nkzB  1okcA  1orsC  1otsA  1p49A  1p4tA  1ppjA
1ppjB  1ppjC  1ppjD  1ppjF  1ppjG  1ppjH  1ppjI  1ppjJ  1qd6C  1qfgA
1qj8A  1qjpA  1qleD  1rzhH  1rzhL  1rzhM  1u19A  1u7gA  1ujwB  1uunA
1uynX  1xioA  1xkwA  1xmeA  1yc9A  1yewA  1yewB  1yewC  1ymgA  1z98A
2a65A  2b2hA  2bhwA  2bl2A  2bs2A  2bs2B  2bs2C  2cfqA  2e74A  2e74B
2e74C  2ei4A  2ervA  2f1vA  2f2bA  2fgqX  2fyuK  2gr8A  2gsmA  2gsmB
2gufA  2h88A  2h88B  2h88C  2h88D  2hdiA  2hdiB  2hydA  2ih3C  2j4uS
2j58A  2j8sA  2jafA  2jlnA  2mprA  2nq2A  2nq2C  2nr9A  2nwlA  2o4vA
2o9gA  2odjA  2porA  2qi9A  2qi9C  2qjyA  2qjyB  2qtkA  2qtsA  2r6gF
2r6gG  2vdfA  2vpzA  2vpzC  2w16A  2w2eA  2wdqA  2wdqB  2wdqC  2wdqD
2wgmA  2wjnC  2wjnH  2wjnL  2wjnM  2wjrA  2wljA  2wswA  2x27X  2x2vA
2x55A  2x9kA  2xfnA  2xovA  2xquA  2y00A  2ydvA  2z73A  2zfgA  2zxeA
2zxeB  2zxeG  3a2sX  3abwA  3aehA  3ag3A  3ag3B  3ag3C  3ag3D  3ag3E
3ag3F  3ag3G  3ag3H  3ag3I  3ag3J  3ag3K  3ag3L  3ag3M  3ar4A  3b9wA
3bs0A  3c02A  3cslA  3cx5A  3cx5B  3cx5C  3cx5D  3cx5F  3cx5G  3cx5H
3cx5I  3d31A  3d9sA  3ddlA  3dh4A  3dwoX  3dzmA  3efmA  3egwA  3egwB
```

```
3egwC  3emnX  3fhhA  3fidA  3gd8A  3giaA  3gp6A  3h90A  3hd6A  3jtyA
3k3fA  3kcuA  3klyA  3kvnA  3l1lA  3ldcA  3m73A  3mp7A  3mp7B  3ne5A
3ne5B  3nsgA  3nymA  3o0rB  3o0rC  3oufA  3pcvA  3pguA  3pikA  3pl9A
3prnA 3qe7A 3qq2A 3rlbA 3rvyA 7ahlA
```

The same was done for 9-mers.

For each sequence, 1000 jobs were run with the following parameters:

```
rosetta_scripts -parser:protocol fnd.xml -database
Rosetta/main/database -in:file:fasta fasta_file -in:file:native
original_pdb -overwrite -use_input_sc -nstruct 100 -jd2:ntrials
10 -mute all -in:file:spanfile span_file -mp:scoring:hbond
-pdb_gz -parser:script_vars frags9mers=9mer_frags_file
-parser:script_vars frags3mers=3mer_frags_file
-parser:script_vars symm_file=denoveo_symm_file
-parser:script_vars span_starts=span_start_position
-parser:script_vars span_ends=span_end_position
-parser:script_vars span_oris=span_orientation
-parser:script_vars span_start_1=span_start_position
-parser:script_vars span_end_1=span_end_position
-parser:script_vars span_start_2=span_start_position
-parser:script_vars span_end_2=span_end_position
-parser:script_vars score_func_0=score0 -parser:script_vars
score_func_1=score1 -parser:script_vars score_func_2=score2
-parser:script_vars score_func_3=score3 -parser:script_vars
score_func_5=score5 -parser:script_vars energy_function=ref
-parser:script_vars steepness=4 -parser:script_vars
membrane_core=10
```

For RosettaMembrane, the steepness is 10, and the membrane core is 15.

The fnd.xml is:

```
<ROSETTASCRIPTS>
  <TASKOPERATIONS>
       <InitializeFromCommandline name="init"/>
       <RestrictToRepacking name="rtr"/>
  </TASKOPERATIONS>
  <SCOREFXNS>
       <ScoreFunction name="score0" weights="%%score_func_0%%" symmetric="1">
       <Reweight scoretype="mp_helicality" weight="100"/>
       </ScoreFunction>
       <ScoreFunction name="score1" weights="%%score_func_1%%" symmetric="1">
       <Reweight scoretype="mp_helicality" weight="100"/>
       </ScoreFunction>
       <ScoreFunction name="score2" weights="%%score_func_2%%" symmetric="1">
       <Reweight scoretype="mp_helicality" weight="100"/>
       </ScoreFunction>
       <ScoreFunction name="score3" weights="%%score_func_3%%" symmetric="1">
```

```
        <Reweight scoretype="mp_helicality" weight="100"/>
        </ScoreFunction>
        <ScoreFunction name="score5" weights="%%score_func_5%%" symmetric="1">
        <Reweight scoretype="mp_helicality" weight="100"/>
        </ScoreFunction>

        <ScoreFunction name="mpframework" weights="mpframework_docking_fa_2015"
symmetric="1"/>
        <ScoreFunction name="mpframeworkNotSymm" weights="mpframework_docking_fa_2015"
symmetric="0"/>
        <ScoreFunction name="ref" weights="ref2015_memb" symmetric="1">
        <Reweight scoretype="mp_helicality" weight="100"/>
        </ScoreFunction>
        <ScoreFunction name="refNotSymm" weights="ref2015_memb" symmetric="0">
        <Reweight scoretype="mp_helicality" weight="100"/>
        </ScoreFunction>

        <ScoreFunction name="helicality" symmetric="1">
        <Reweight scoretype="mp_helicality" weight="1"/>
        </ScoreFunction>
        <ScoreFunction name="helicality_notsymm" symmetric="0">
        <Reweight scoretype="mp_helicality" weight="1"/>
        </ScoreFunction>
   </SCOREFXNS>
<RESIDUE_SELECTORS>
   <Layer name="layer" select_core="1" select_boundary="1" select_surface="1"/>
 </RESIDUE_SELECTORS>
   <MOVERS>
        <SetupForSymmetry name="symm" definition="%%symm_file%%"/>
        <SymmetricAddMembraneMover name="add_memb" membrane_core="%%membrane_core%%"
steepness="%%steepness%%" span_starts_num="%%span_starts%%"
span_ends_num="%%span_ends%%" span_orientations="%%span_oris%%"/>
        <MembranePositionFromTopologyMover name="init_pos"/>
        <FastRelax name="fast_relax" scorefxn="%%energy_function%%"
task_operations="init"/>

        Fragment movers
        <SingleFragmentMover name="frag9" fragments="%%frags9mers%%" policy="uniform">
        <MoveMap>
        <Span begin="1" end="24" chi="1" bb="1"/>
        </MoveMap>
        </SingleFragmentMover>
        <SingleFragmentMover name="frag3" fragments="%%frags3mers%%" policy="smooth">
        <MoveMap>
        <Span begin="1" end="24" chi="1" bb="1"/>
        </MoveMap>
        </SingleFragmentMover>

        Fold-and-dock specific movers
        <SymFoldandDockRbTrialMover name="rbtrial" rot_mag="8.0" trans_mag="3.0"
rotate_anchor_to_x="1"/>
        <SymFoldandDockRbTrialMover name="rbtrial_smooth" rot_mag="1.0" trans_mag="0.1"
rotate_anchor_to_x="1"/>
        <SymFoldandDockMoveRbJumpMover name="rbjump"/>
        <SymFoldandDockSlideTrialMover name="slidetrial"/>

        Random movers
        <RandomMover name="early_stage_moveset"
movers="frag9,rbtrial,rbjump,slidetrial" weights="1.0,0.2,1.0,0.1" repeats="1"/>
```

```
       <RandomMover name="final_stage_moveset"
movers="frag3,rbtrial_smooth,rbjump,slidetrial" weights="1.0,0.2,1.0,0.1"
repeats="1"/>

       Monte Carlo Movers
       <GenericMonteCarlo name="stage1" scorefxn_name="score0"
mover_name="early_stage_moveset" temperature="2.0" trials="200" recover_low="1"/>
       <GenericMonteCarlo name="stage2" scorefxn_name="score1"
mover_name="early_stage_moveset" temperature="2.0" trials="200" recover_low="1"/>
       <GenericMonteCarlo name="stage3a" scorefxn_name="score2"
mover_name="early_stage_moveset" temperature="2.0" trials="20" recover_low="1"/>
       <GenericMonteCarlo name="stage3b" scorefxn_name="score5"
mover_name="early_stage_moveset" temperature="2.0" trials="20" recover_low="1"/>
       <GenericMonteCarlo name="stage4" scorefxn_name="score3"
mover_name="final_stage_moveset" temperature="2.0" trials="400" recover_low="1"/>

       Special stage 3 logic
       <ParsedProtocol name="stage3_cyc">
       <Add mover="stage3a"/>
       <Add mover="stage3b"/>
       </ParsedProtocol>
       <LoopOver name="stage3" mover_name="stage3_cyc" iterations="5" drift="1"/>

       Converts the centroid-level pose to fullatom for scoring
       <SwitchResidueTypeSetMover name="fullatom" set="fa_standard"/>
       <ExtractAsymmetricPose name="extract_asp" clear_sym_def="1"/>
       <MinMover name="min_mover" scorefxn="refNotSymm" chi="1" bb="1" jump="1"/>
       <PackRotamersMover name="pack" scorefxn="refNotSymm"
task_operations="init,rtr"/>
       <RotamerTrialsMinMover name="RTmin" scorefxn="refNotSymm"
task_operations="init,rtr"/>
       <DumpPdb name="dump_pdb" fname="dump.pdb" scorefxn="%%energy_function%%"/>
  </MOVERS>
  <FILTERS>
       <ScoreType name="total" scorefxn="%%energy_function%%" score_type="total_score"
confidence="1" threshold="0"/>
       <Sasa name="a_sasa" confidence="1" threshold="300"/>

       <ResidueLipophilicity name="a_res_lipo" threshold="1000" confidence="0"/>
       <SpanTopologyMatchPose name="a_span_topo" confidence="0"/>
       <Ddg name="a_ddg" scorefxn="%%energy_function%%NotSymm" chain_num="2"
repeats="5" extreme_value_removal="true" confidence="1" threshold="-5"/>
       <PackStat name="a_pack" confidence="1" threshold="0.3"/>
       <BuriedUnsatHbonds2 name="a_unsat" scorefxn="%%energy_function%%"
confidence="0"/>
       <ShapeComplementarity name="a_shape" confidence="0"/>
       <TMsSpanMembrane name="a_tms_span" confidence="1" min_distance="25"/>
       <TMsSpanMembrane name="a_tms_span_fa" confidence="1" min_distance="25"/>
   <HelixHelixAngle name="a_hha_ang" angle_or_dist="angle"
start_helix_1="%%span_start_1%%" end_helix_1="%%span_end_1%%"
start_helix_2="%%span_start_2%%" end_helix_2="%%span_end_2%%" confidence="0"/>
       <HelixHelixAngle name="a_hha_dst_vec" angle_or_dist="dist" dist_by_atom="0"
start_helix_1="%%span_start_1%%" end_helix_1="%%span_end_1%%"
start_helix_2="%%span_start_2%%" end_helix_2="%%span_end_2%%" confidence="0"/>
       <HelixHelixAngle name="a_hha_dst_atm" angle_or_dist="dist" dist_by_atom="1"
start_helix_1="%%span_start_1%%" end_helix_1="%%span_end_1%%"
start_helix_2="%%span_start_2%%" end_helix_2="%%span_end_2%%" confidence="0"/>
       <MembAccesResidueLipophilicity name="a_marl" confidence="0" verbose="0"/>
       <ScoreType name="a_helicality" scorefxn="helicality_notsymm"
score_type="mp_helicality" confidence="1" threshold="10"/>
```

```
      <ScoreType name="a_helicality_symm" scorefxn="helicality"
score_type="mp_helicality" confidence="1" threshold="10"/>
      <MPSpanAngle name="a_angle_1" tm="1" ang_min="0" ang_max="50" confidence="1"/>
      <MPSpanAngle name="a_angle_2" tm="2" ang_min="0" ang_max="50" confidence="1"/>
      <RmsdFromResidueSelector name="a_rmsd" CA_only="1" reference_selector="layer"
query_selector="layer" confidence="1" threshold="15"/>
      <BindingStrain name="a_bind" scorefxn="%%energy_function%%NotSymm" jump="1"
confidence="1" threshold="5"/>
      <PoseInfo name="info"/>
  </FILTERS>
  <PROTOCOLS>
   <Add mover="symm"/>
   <Add mover="add_memb"/>

   <Add mover="stage1"/>
   <Add mover="stage2"/>
   <Add mover="stage3"/>
   <Add mover="stage4"/>

   <Add filter="a_helicality_symm"/>
   <Add filter="a_angle_1"/>
   <Add filter="a_angle_2"/>

   <Add mover="fullatom"/>

   <Add filter="a_tms_span"/>
   <Add mover="fast_relax"/>

   <Add filter="total"/>
   <Add filter="a_sasa"/>

   <Add filter="a_span_topo"/>

   <Add mover="extract_asp"/>
   <Add mover="pack"/>
   <Add mover="min_mover"/>
   <Add mover="RTmin"/>
   <Add mover="RTmin"/>

   <Add filter="a_tms_span"/>
   <Add filter="total"/>
   <Add filter="a_sasa"/>
   <Add filter="a_span_topo"/>

   <Add filter="a_rmsd"/>
   <Add filter="a_res_lipo"/>
   <Add filter="a_pack"/>
   <Add filter="a_unsat"/>
   <Add filter="a_shape"/>
  <Add filter="a_ddg"/>
   <Add filter="a_hha_ang"/>
   <Add filter="a_hha_dst_vec"/>
   <Add filter="a_hha_dst_atm"/>
   <Add filter="a_marl"/>
   <Add filter="a_tms_span_fa"/>
   <Add filter="a_helicality"/>
   <Add filter="a_angle_1"/>
   <Add filter="a_angle_2"/>
   <Add filter="a_bind"/>
  </PROTOCOLS>
  <OUTPUT scorefxn="%%energy_function%%NotSymm"/>
```

```
</ROSETTASCRIPTS>
```

1000 instances of the above command were executed, creating up to 100,000 models. Models were filtered using the following criteria: score in the bottom 10%, SASA > 500 Å$^2$, shape complementarity > 0.6, $\Delta\Delta G_{binding}$ < -5 R.e.u., binding strain < 4 R.e.u. and mp_helicality < 0.1 R.e.u. For homodimers, the distance between the closest atoms on the helices was filtered to be < 9 Å, as calculated by the filter HelixHelixAngle.

The filtered models were then score-wise clustered. Iteratively, all models were aligned to the best scoring model, and ones closer than 4 Å were removed. Alignment and RMSD calculations were computed in PyMOL. And the five largest clusters are reported using the cluster representative with the lowest energy.

As a first step, each structure was refined, using an either asymmetric or symmetric protocol, as appropriate.

Refinement command line:

```
~/Rosetta/main/source/bin/rosetta_scripts.default.linuxgccreleas
e -parser:protocol refine.xml -s PDB_FILE -overwrite
-script_vars cst_value=0.4 -script_vars
cst_full_path=COORD_CST_PATH -script_vars
symm_file=SYMM_FILE_PATH -extrachi_cutoff 10
-ignore_unrecognized_res -chemical:exclude_patches LowerDNA
UpperDNA Cterm_amidation SpecialRotamer VirtualBB ShoveBB
VirtualDNAPhosphate VirtualNTerm CTermConnect sc_orbitals
pro_hydroxylated_case1 pro_hydroxylated_case2 ser_phosphorylated
thr_phosphorylated  tyr_phosphorylated tyr_sulfated
lys_dimethylated lys_monomethylated  lys_trimethylated
lys_acetylated glu_carboxylated cys_acetylated tyr_diiodinated
N_acetylated C_methylamidated MethylatedProteinCterm
-script_vars span_starts=COMMA_SEPARATED_SPAN_START_POSITIONS
-script_vars span_ends=COMMA_SEPARATED_SPAN_END_POSITIONS
-script_vars span_oris=COMMA_SEPARATED_ORIENTATIONS
-parser:script_vars membrane_core=MEMBRANE_CORE
-parser:script_vars steepness=STEEPNESS -script_vars mpf=MPF
-mp:scoring:hbond
```

Where SYMM_FILE is generated by the ~/Rosetta/main/source//src/apps/public/symmetry//make_symmdef_file.pl script from the

Rosetta modelling suite. COORD_CST_FILE is a list of coordinate constraints for all atoms in the structure. MEMBRANE_CORE is 15 for RosettaMembrane, 10 for ref2015_memb, and irrelevant for ref2015. STEEPNESS is 10 for RosettaMembrane, 4 for ref2015_memb, and irrelevant for ref2015. The mpf script variable is used to differentiate functions and is only used for RosettaMembrane, with the value _mpf.

RosettaScripts protocol for asymmetric refinement using RosettaMembrane or ref2015_memb:

```
<ROSETTASCRIPTS>
  <SCOREFXNS>
    <ScoreFunction name="full" weights="ref2015_memb" symmetric="0">
      <Reweight scoretype="coordinate_constraint" weight="%%cst_value%%"/>
    </ScoreFunction>
    <ScoreFunction name="soft" weights="ref2015_soft" symmetric="0">
      <Reweight scoretype="mp_res_lipo" weight="1"/>
      <Reweight scoretype="coordinate_constraint" weight="%%cst_value%%"/>
    </ScoreFunction>
    <ScoreFunction name="ref_pure" weights="ref2015_memb" symmetric="0"/>
    <ScoreFunction name="helicality" symmetric="1">
      <Reweight scoretype="mp_helicality" weight="1"/>
    </ScoreFunction>

    <ScoreFunction name="full_mpf" weights="mpframework_docking_fa_2015"
symmetric="0">
      <Reweight scoretype="coordinate_constraint" weight="%%cst_value%%"/>
    </ScoreFunction>
    <ScoreFunction name="soft_mpf" weights="mpframework_docking_fa_2015"
symmetric="0">
      <Reweight scoretype="coordinate_constraint" weight="%%cst_value%%"/>
    </ScoreFunction>
    <ScoreFunction name="ref_pure_mpf" weights="mpframework_docking_fa_2015.wts"
symmetric="0"/>
  </SCOREFXNS>
  <RESIDUE_SELECTORS>
  </RESIDUE_SELECTORS>
  <TASKOPERATIONS>
    <InitializeFromCommandline name="init"/>
    <RestrictToRepacking name="rtr"/>
  </TASKOPERATIONS>
  <MOVERS>
    <AddMembraneMover name="add_memb" membrane_core="10" steepness="4"
span_starts="%%span_starts%%" span_ends="%%span_ends%%"
span_orientations="%%span_oris%%"/>
    <PackRotamersMover name="soft_repack" scorefxn="soft%%mpf%%"
task_operations="init,rtr"/>
    <PackRotamersMover name="hard_repack" scorefxn="full%%mpf%%"
task_operations="init,rtr"/>
    <RotamerTrialsMinMover name="RTmin" scorefxn="full" task_operations="init,rtr"/>
    <MinMover name="soft_min" scorefxn="soft%%mpf%%" chi="1" bb="1" jump="0"/>
    <MinMover name="hard_min" scorefxn="full%%mpf%%" chi="1" bb="1" jump="0"/>
    <ConstraintSetMover name="add_CA_cst" cst_file="%%cst_full_path%%"/>
    <ParsedProtocol name="refinement_block"> #10 movers
      <Add mover_name="soft_repack"/>
      <Add mover_name="soft_min"/>
      <Add mover_name="soft_repack"/>
```

```
      <Add mover_name="hard_min"/>
      <Add mover_name="hard_repack"/>
      <Add mover_name="hard_min"/>
      <Add mover_name="hard_repack"/>
      <Add mover_name="RTmin"/>
      <Add mover_name="RTmin"/>
      <Add mover_name="hard_min"/>
    </ParsedProtocol>
    <LoopOver name="iter4" mover_name="refinement_block" iterations="4"/> #16
reacpk+min iterations total
    <DumpPdb name="dump_pdb" fname="dump.pdb"/>
  </MOVERS>
  <FILTERS>
    <ScoreType name="stability_score_full" scorefxn="full%%mpf%%"
score_type="total_score" confidence="0" threshold="0"/>
    <ScoreType name="stability_pure" scorefxn="ref_pure%%mpf%%"
score_type="total_score" confidence="0" threshold="0"/>
    <Rmsd name="rmsd" confidence="0"/>
    <ResidueLipophilicity name="a_res_lipo" threshold="1000" confidence="0"/>
    <SpanTopologyMatchPose name="a_span_topo" confidence="0"/>
    <TMsSpanMembrane name="a_tms_span" confidence="0" min_distance="25"/>
    <MembAccesResidueLipophilicity name="a_marl" confidence="0" verbose="0"/>
    <ScoreType name="a_helicality" scorefxn="helicality" score_type="mp_helicality"
confidence="0" threshold="10"/>
    <Time name="timer"/>
  </FILTERS>
  <PROTOCOLS>
    <Add mover="add_memb"/>
    <Add filter="timer"/>
    <Add mover="add_CA_cst"/>
    <Add mover="iter4"/>
    <Add filter="stability_score_full"/>
    <Add filter="stability_pure"/>
    <Add filter="a_res_lipo"/>
    <Add filter="a_span_topo"/>
    <Add filter="a_tms_span"/>
    <Add filter="a_marl"/>
    <Add filter="a_helicality"/>
    <Add filter="timer"/>
  </PROTOCOLS>
  <OUTPUT scorefxn="full%%mpf%%"/>
</ROSETTASCRIPTS>
```

RosettaScripts protocol for asymmetric refinement using ref2015:

```
<ROSETTASCRIPTS>
  <SCOREFXNS>
    <ScoreFunction name="full" weights="ref2015" symmetric="0">
      <Reweight scoretype="coordinate_constraint" weight="%%cst_value%%"/>
    </ScoreFunction>
    <ScoreFunction name="soft" weights="ref2015_soft" symmetric="0">
      <Reweight scoretype="coordinate_constraint" weight="%%cst_value%%"/>
    </ScoreFunction>
    <ScoreFunction name="ref_pure" weights="ref2015" symmetric="0"/>
  </SCOREFXNS>
  <RESIDUE_SELECTORS>
  </RESIDUE_SELECTORS>
  <TASKOPERATIONS>
    <InitializeFromCommandline name="init"/>
    <RestrictToRepacking name="rtr"/>
```

```
  </TASKOPERATIONS>
  <MOVERS>
    <PackRotamersMover name="soft_repack" scorefxn="soft%%mpf%%"
task_operations="init,rtr"/>
    <PackRotamersMover name="hard_repack" scorefxn="full%%mpf%%"
task_operations="init,rtr"/>
    <RotamerTrialsMinMover name="RTmin" scorefxn="full" task_operations="init,rtr"/>
    <MinMover name="soft_min" scorefxn="soft%%mpf%%" chi="1" bb="1" jump="0"/>
    <MinMover name="hard_min" scorefxn="full%%mpf%%" chi="1" bb="1" jump="0"/>
    <ConstraintSetMover name="add_CA_cst" cst_file="%%cst_full_path%%"/>
    <ParsedProtocol name="refinement_block"> #10 movers
      <Add mover_name="soft_repack"/>
      <Add mover_name="soft_min"/>
      <Add mover_name="soft_repack"/>
      <Add mover_name="hard_min"/>
      <Add mover_name="hard_repack"/>
      <Add mover_name="hard_min"/>
      <Add mover_name="hard_repack"/>
      <Add mover_name="RTmin"/>
      <Add mover_name="RTmin"/>
      <Add mover_name="hard_min"/>
    </ParsedProtocol>
    <LoopOver name="iter4" mover_name="refinement_block" iterations="4"/> #16
reacpk+min iterations total
    <DumpPdb name="dump_pdb" fname="dump.pdb"/>
  </MOVERS>
  <FILTERS>
    <ScoreType name="stability_score_full" scorefxn="full%%mpf%%"
score_type="total_score" confidence="0" threshold="0"/>
    <ScoreType name="stability_pure" scorefxn="ref_pure%%mpf%%"
score_type="total_score" confidence="0" threshold="0"/>
    <Rmsd name="rmsd" confidence="0"/>
    <Time name="timer"/>
  </FILTERS>
  <PROTOCOLS>
    <Add filter="timer"/>
    <Add mover="add_CA_cst"/>
    <Add mover="iter4"/>
    <Add filter="stability_score_full"/>
    <Add filter="stability_pure"/>
    <Add filter="timer"/>
  </PROTOCOLS>
  <OUTPUT scorefxn="full%%mpf%%"/>
</ROSETTASCRIPTS>
```

RosettaScripts protocol for symmetric refinement using RosettaMembrane or ref2015_memb:

```
<ROSETTASCRIPTS>
  <TASKOPERATIONS>
    <InitializeFromCommandline name="init"/>
    <RestrictToRepacking name="rtr"/>
  </TASKOPERATIONS>
  <SCOREFXNS>
    <ScoreFunction name="full" weights="ref2015_memb" symmetric="1">
      <Reweight scoretype="coordinate_constraint" weight="%%cst_value%%"/>
    </ScoreFunction>
    <ScoreFunction name="soft" weights="ref2015_soft" symmetric="1">
      <Reweight scoretype="mp_res_lipo" weight="1"/>
      <Reweight scoretype="coordinate_constraint" weight="%%cst_value%%"/>
    </ScoreFunction>
```

```
    <ScoreFunction name="ref_pure" weights="ref2015_memb" symmetric="1"/>
    <ScoreFunction name="helicality" symmetric="1">
      <Reweight scoretype="mp_helicality" weight="1"/>
    </ScoreFunction>

    <ScoreFunction name="full_mpf" weights="mpframework_docking_fa_2015"
symmetric="1">
      <Reweight scoretype="coordinate_constraint" weight="%%cst_value%%"/>
    </ScoreFunction>
    <ScoreFunction name="soft_mpf" weights="mpframework_docking_fa_2015"
symmetric="1">
      <Reweight scoretype="coordinate_constraint" weight="%%cst_value%%"/>
    </ScoreFunction>
    <ScoreFunction name="ref_pure_mpf" weights="mpframework_docking_fa_2015"
symmetric="1"/>
  </SCOREFXNS>
  <RESIDUE_SELECTORS>
  </RESIDUE_SELECTORS>
  <MOVERS>
    <SymmetricAddMembraneMover name="add_memb" membrane_core="10" steepness="4"
span_starts="%%span_starts%%" span_ends="%%span_ends%%"
span_orientations="%%span_oris%%"/>
    <SetupForSymmetry name="symm" definition="%%symm_file%%"/>
    <SymPackRotamersMover name="soft_repack" scorefxn="soft%%mpf%%"
task_operations="init,rtr"/>
    <SymPackRotamersMover name="hard_repack" scorefxn="full%%mpf%%"
task_operations="init,rtr"/>
    <RotamerTrialsMinMover name="RTmin" scorefxn="full" task_operations="init,rtr"/>
    <SymMinMover name="soft_min" scorefxn="soft%%mpf%%" chi="1" bb="1" jump="0"/>
    <SymMinMover name="hard_min" scorefxn="full%%mpf%%" chi="1" bb="1" jump="0"/>
    <ConstraintSetMover name="add_CA_cst" cst_file="%%cst_full_path%%"/>
    <ParsedProtocol name="refinement_block"> #10 movers
      <Add mover_name="soft_repack"/>
      <Add mover_name="soft_min"/>
      <Add mover_name="soft_repack"/>
      <Add mover_name="hard_min"/>
      <Add mover_name="hard_repack"/>
      <Add mover_name="hard_min"/>
      <Add mover_name="hard_repack"/>
      <Add mover_name="RTmin"/>
      <Add mover_name="RTmin"/>
      <Add mover_name="hard_min"/>
    </ParsedProtocol>
    <LoopOver name="iter4" mover_name="refinement_block" iterations="4"/> #16
reacpk+min iterations total
    <DumpPdb name="dump_pdb" fname="dump.pdb"/>
  </MOVERS>
  <FILTERS>
    <ScoreType name="stability_score_full" scorefxn="full%%mpf%%"
score_type="total_score" confidence="0" threshold="0"/>
    <ScoreType name="stability_pure" scorefxn="ref_pure%%mpf%%"
score_type="total_score" confidence="0" threshold="0"/>
    <Rmsd name="rmsd" confidence="0"/>
    <ResidueLipophilicity name="a_res_lipo" threshold="1000" confidence="0"/>
    <SpanTopologyMatchPose name="a_span_topo" confidence="0"/>
    <TMsSpanMembrane name="a_tms_span" confidence="0" min_distance="25"/>
    <MembAccesResidueLipophilicity name="a_marl" confidence="0" verbose="0"/>
    <ScoreType name="a_helicality" scorefxn="helicality" score_type="mp_helicality"
confidence="0" threshold="10"/>
    <Time name="timer"/>
  </FILTERS>
```

```
  <PROTOCOLS>
    <Add mover="symm"/>
    <Add mover="add_memb"/>
    <Add filter="timer"/>
    <Add mover="add_CA_cst"/>
    <Add mover="iter4"/>
    <Add filter="stability_score_full"/>
    <Add filter="stability_pure"/>
    <Add filter="a_res_lipo"/>
    <Add filter="a_span_topo"/>
    <Add filter="a_tms_span"/>
    <Add filter="a_marl"/>
    <Add filter="a_helicality"/>
    <Add filter="timer"/>
  </PROTOCOLS>
  <OUTPUT scorefxn="full%%mpf%%"/>
</ROSETTASCRIPTS>
```

RosettaScripts protocol for symmetric refinement using ref2015:

```
<ROSETTASCRIPTS>
  <TASKOPERATIONS>
    <InitializeFromCommandline name="init"/>
    <RestrictToRepacking name="rtr"/>
  </TASKOPERATIONS>
  <SCOREFXNS>
    <ScoreFunction name="full" weights="ref2015" symmetric="1">
      <Reweight scoretype="coordinate_constraint" weight="%%cst_value%%"/>
    </ScoreFunction>
    <ScoreFunction name="soft" weights="ref2015_soft" symmetric="1">
      <Reweight scoretype="coordinate_constraint" weight="%%cst_value%%"/>
    </ScoreFunction>
    <ScoreFunction name="ref_pure" weights="ref2015" symmetric="1"/>
  </SCOREFXNS>
  <RESIDUE_SELECTORS>
  </RESIDUE_SELECTORS>
  <MOVERS>
    <SetupForSymmetry name="symm" definition="%%symm_file%%"/>
    <SymPackRotamersMover name="soft_repack" scorefxn="soft%%mpf%%"
task_operations="init,rtr"/>
    <SymPackRotamersMover name="hard_repack" scorefxn="full%%mpf%%"
task_operations="init,rtr"/>
    <RotamerTrialsMinMover name="RTmin" scorefxn="full" task_operations="init,rtr"/>
    <SymMinMover name="soft_min" scorefxn="soft%%mpf%%" chi="1" bb="1" jump="0"/>
    <SymMinMover name="hard_min" scorefxn="full%%mpf%%" chi="1" bb="1" jump="0"/>
    <ConstraintSetMover name="add_CA_cst" cst_file="%%cst_full_path%%"/>
    <ParsedProtocol name="refinement_block"> #10 movers
      <Add mover_name="soft_repack"/>
      <Add mover_name="soft_min"/>
      <Add mover_name="soft_repack"/>
      <Add mover_name="hard_min"/>
      <Add mover_name="hard_repack"/>
      <Add mover_name="hard_min"/>
      <Add mover_name="hard_repack"/>
      <Add mover_name="RTmin"/>
      <Add mover_name="RTmin"/>
      <Add mover_name="hard_min"/>
    </ParsedProtocol>
    <LoopOver name="iter4" mover_name="refinement_block" iterations="4"/> #16
reacpk+min iterations total
```

```
    <DumpPdb name="dump_pdb" fname="dump.pdb"/>
  </MOVERS>
  <FILTERS>
    <ScoreType name="stability_score_full" scorefxn="full%%mpf%%"
score_type="total_score" confidence="0" threshold="0"/>
    <ScoreType name="stability_pure" scorefxn="ref_pure%%mpf%%"
score_type="total_score" confidence="0" threshold="0"/>
    <Rmsd name="rmsd" confidence="0"/>
    <Time name="timer"/>
  </FILTERS>
  <PROTOCOLS>
    <Add mover="symm"/>
    <Add filter="timer"/>
    <Add mover="add_CA_cst"/>
    <Add mover="iter4"/>
    <Add filter="stability_score_full"/>
    <Add filter="stability_pure"/>
    <Add filter="timer"/>
  </PROTOCOLS>
  <OUTPUT scorefxn="full%%mpf%%"/>
</ROSETTASCRIPTS>
```

For each structure, 10 trajectories were attempted, using each energy function. The best scoring model was used for the design step. Symmetry and coordinate constraints files were recreated to fit the refined model.

Design for sequence recovery benchmark:

The following command line was used for each structure:

```
~/Rosetta/main/source/bin/rosetta_scripts.default.linuxgccreleas
e -parser:protocol design.xml -s BEST_MODEL_PATH -overwrite
-script_vars scfxn=SCORE_FUNCATION_NAME -script_vars
symm_file=BEST_MODEL_SYMM_FILE -extrachi_cutoff 10
-ignore_unrecognized_res -chemical:exclude_patches LowerDNA
UpperDNA Cterm_amidation SpecialRotamer VirtualBB ShoveBB
VirtualDNAPhosphate VirtualNTerm CTermConnect sc_orbitals
pro_hydroxylated_case1 pro_hydroxylated_case2 ser_phosphorylated
thr_phosphorylated  tyr_phosphorylated tyr_sulfated
lys_dimethylated lys_monomethylated  lys_trimethylated
lys_acetylated glu_carboxylated cys_acetylated tyr_diiodinated
N_acetylated C_methylamidated MethylatedProteinCterm
-script_vars span_starts=COMMA_SEPARATED_SPAN_START_POSITIONS
-script_vars span_ends=COMMA_SEPARATED_SPAN_ENDS_POSITIONS
-script_vars
span_oris=COMMA_SEPARATED_SPAN_ORIENTATION_POSITIONS
-parser:script_vars membrane_core=MEMBRANE_CORE
-parser:script_vars steepness=STEEPNESS
-mp:scoring:hbond -script_vars add_memb=ADD_MEMB
```

Where SCORE_FUNCTION_PATH is either mpframework_docking_fa_2015 for RosettaMembrane, ref2015_memb or ref2015. Other attributes are as described for refinement.

RosettaScripts protocol for asymmetric design:

```
<ROSETTASCRIPTS>
  <SCOREFXNS>
    <ScoreFunction name="full" weights="%%scfxn%%" symmetric="0">
    </ScoreFunction>
  </SCOREFXNS>
  <RESIDUE_SELECTORS>
  </RESIDUE_SELECTORS>
  <TASKOPERATIONS>
    <InitializeFromCommandline name="init"/>
  </TASKOPERATIONS>
  <MOVERS>
    %%add_memb%%AddMembraneMover name="add_memb" membrane_core="%%membrane_core%%"
steepness="%%steepness%%" span_starts="%%span_starts%%" span_ends="%%span_ends%%"
span_orientations="%%span_oris%%"/>
    <PackRotamersMover name="repack" scorefxn="full" task_operations="init"/>
  </MOVERS>
  <FILTERS>
    <ScoreType name="stability_score_full" scorefxn="full" score_type="total_score"
confidence="0" threshold="0"/>
    <Time name="timer"/>
  </FILTERS>
  <PROTOCOLS>
    <Add filter="timer"/>
    %%add_memb%%Add mover="add_memb"/>
    <Add mover="repack"/>
    <Add filter="stability_score_full"/>
    <Add filter="timer"/>
  </PROTOCOLS>
  <OUTPUT scorefxn="full"/>
</ROSETTASCRIPTS>
```

RosettaScripts protocol for symmetric design:

```
<ROSETTASCRIPTS>
  <SCOREFXNS>
    <ScoreFunction name="full" weights="%%scfxn%%" symmetric="1">
    </ScoreFunction>
  </SCOREFXNS>
  <RESIDUE_SELECTORS>
  </RESIDUE_SELECTORS>
  <TASKOPERATIONS>
    <InitializeFromCommandline name="init"/>
  </TASKOPERATIONS>
  <MOVERS>
    %%add_memb%%SymmetricAddMembraneMover name="add_memb" membrane_core="10"
steepness="4" span_starts="%%span_starts%%" span_ends="%%span_ends%%"
span_orientations="%%span_oris%%"/>
    <SetupForSymmetry name="symm" definition="%%symm_file%%"/>
    <SymPackRotamersMover name="repack" scorefxn="full" task_operations="init"/>
  </MOVERS>
```

```
  <FILTERS>
    <ScoreType name="stability_score_full" scorefxn="full" score_type="total_score"
confidence="0" threshold="0"/>
    <Time name="timer"/>
  </FILTERS>
  <PROTOCOLS>
    <Add filter="timer"/>
    <Add mover="symm"/>
    %%add_memb%%Add mover="add_memb"/>
    <Add mover="repack"/>
    <Add filter="stability_score_full"/>
    <Add filter="timer"/>
  </PROTOCOLS>
  <OUTPUT scorefxn="full"/>
</ROSETTASCRIPTS>
```
